# Study on Linear Combination of Long Memory Processes Corrupted by Additive Noises for fMRI Time Series Analysis

**DOI:** 10.1101/210518

**Authors:** Wonsang You, Catherine Limperopoulos

## Abstract

Estimating the long memory parameter of the fMRI time series enables us to understand the fractal behavior of neural activity of the brain through fMRI time series. However, the existence of white noise and physiological noise compounds which also have fractal properties prevent us from making the estimation precise. As basic strategies to overcome noises, we address how to estimate the long memory parameter in the presence of additive noises, and how to estimate the long memory parameters of linearly combined long memory processes.

## 1 Introduction

Fractal can be regarded as self-similarity of an object which means that the statistical properties remain invariant throughout different scales. It is well known that many natural phenomena have such a fractal structure; for example, it can be found at river networks, clouds, trees, neuronal networks, Koch curve, and so forth [11]. The fMRI time series have also fractal properties or 1/*f* spectral densities [3][12]. Thus, some neuroscientists have attempted to study fractal properties of fMRI time series by estimating the fractal dimension or long memory parameter. In results, they have observed not only that the fMRI time series tend to be corrupted by a fractional Gaussian noise (fGn) which has fractal properties, but also that the Hurst exponent of the fGn noise provides valuable information on Alzheimer’s disease [12], Moreover, the fractal properties are stronger in grey matter which has higher population of neurons, than in white matter of the brain even during resting state [4]. These reports indicate that the fractal properties of fMRI time series may represent neural activity of the brain. Thus, it is manifest that the fractal properties are one of important features to analyze fMRI time series.

Our interest is to estimate the long memory parameter of the endogenous signal which can be modeled as a long memory process in order to track the spontaneous neural activity of the brain during resting state of a subject through analyzing fMRI time series. Unfortunately, the data normally tend to suffer from tremendous noise compounds such as cardiac pulse, respiration, subject motion, and scanner noise. While it is not serious in task-based experiments, it could be a critical problem in resting state fMRI analysis since it may significantly prevent us from extracting the endogenous signal which arises from neural activity of the brain. It means that the long memory behavior of a raw fMRI time series may be attributed not only to endogenous neural activity but also to physiological noise, subject movement, and other sources [13][14]. To precisely estimate the long memory parameter of a pure endogenous signal, we should model the raw fMRI time series as the linear sum of several long memory processes (for example, the summation of endogenous signal and physiological pulsations) and additive noises. Under this model, our works consist of two steps: (1) estimating the parameters of each component in the model, and (2) classifying each component into endogenous signal, physiological noises, or other additive noises.

Here, we focus on the first step: estimating the parameters of linearly combined long memory processes in the presence of additive noises. To be specific, we address two topics; one is the parameter estimation of a long memory process in the presence of additive noises, and the other is the parameter estimation of linearly combined long memory processes. Regarding the former topic, we thought that Achard’s method for a fractional Brownian motion corrupted by additive noises [7] can be also applicable to estimate the parameter of a long memory process corrupted by additive noise. In the latter topic, we will introduce a method to estimate the parameters of linearly combined long memory processes. The fact that the linear combination of long memory processes results in the linear combination of wavelet variances allows us to use multiexponential analysis in order to estimate the parameters of linearly combined long memory processes.

## 2 The Signal Model

Let **Y** := {**Y**_*l*_}_l=1,⋯,*L*_ be a set of *L* stochastic processes where **Y**_*l*_ := {*Y_l_*(*t*)}_*t*=1,⋯, *N*_ for *l* =1, ⋯, *L*. In other words, we have *L* fMRI time series with *N* time points. We assume that each process can be modeled as the linear combination of endogenous signal, cardiac noise, respiratory noise, and white noise as follows

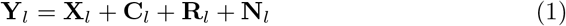

where **X**_*l*_ := {*X_l_*(*t*)}_*t*=1,⋯, *N*_ is an endogenous signal at *l*-th voxel, **C**_*l*_ := {*C_l_*(*t*)}_*t*=1,⋯, *N*_ is a cardiac noise at *l*-th voxel, **R**_*l*_ := {*R_l_*(*t*)}_*t*=1,⋯, *N*_ is a respiratory noise at *l*-th voxel, and **N**_*l*_ := {*N_l_*(*t*)}_*t*=1,⋯, *N*_ is an additive white Gaussian noise at *l*-th voxel. That is, *N_l_*(*t*) for *t* = 1, ⋯, *N* are *i.i.d*. Gaussian random variable with variance 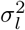. Since the first step is to eliminate the additive white noise **N**_*l*_, we can simplify the above equation into

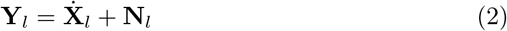

where 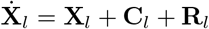.

## 3 Parameter Estimation of a Long Memory Process Corrupted by Additive Noises

### 3.1 Some Definitions

As we discussed in the previous section, the endogenous signal **X**_*l*_ can be modeled as a type of long memory process. Likewise, we can also regard cardiac noise C; and respiratory noise **R**_*l*_ as long memory processes. Indeed, there have been a lot of evidences which imply that heartbeat [16][17][18][19][20][21] and respiration [22][23][24] also have fractal properties.

We suppose that the linear summation of long memory processes is approximately a long memory process. In other words, if each process *X_k_* for *k* = 1, ⋯, *N* has long memory, the process

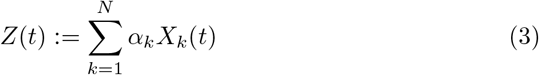

also has long memory for 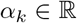. We did not prove this hypothesis but we will indirectly see at the section 4 how reasonable it is. In addition, Christoph Thäle [25] proved that the linear combination of *N* fractional Brownian motions with *H_k_* > 1/2 for *k* =1, ⋯, *N* is also long-range dependent. Under this hypothesis, 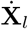 in (2) can also be regarded as a long memory process.

We adopt the definition of a long memory process suggested by Moulines *et al*. [5]; that is, a real-valued discrete process 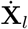 is said to have memory parameter *d_l_* (and is called a *M*(*d_l_*) process) if, for any integer *D* > *d_l_* – 1/2, the *D*-th order difference process 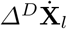 (where 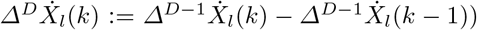) is weakly stationary with spectral density function

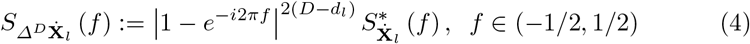

where 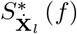 is a non-negative symmetric function which is bounded on (−1/2,1/2).

To estimate the long memory parameter *d_l_*, let us define a filter *a* of length *p* +1 satisfying the following conditions^1^:

1. 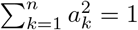.
2. There exists *α* > 0 such that 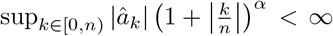 where 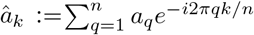.
3. It has *M* vanishing moments, i.e. 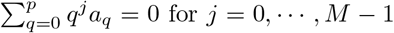, and 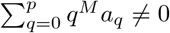.

On the other hand, the dilated filter^2^ *a_j_* is defined as the filter of length 2*^j^p* + 1 such that for *j* ≥ 0 and *k* = 0, ⋯, 2*^j^p*,

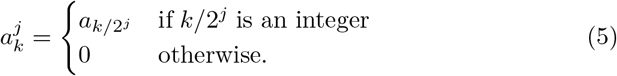

Let 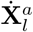 be the vector 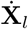 filtered with *a* such that for *i* = *l* + 1, ⋯, *n*

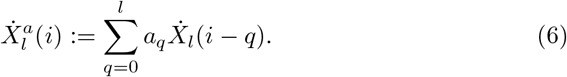

### 3.2 Asymptotic Approximation of Spectral Density

Moulines *et al*. showed that the spectral density of the wavelet coefficients of an *M*(*d*) process can be asymptotically approximated by that of fractional Brownian motion (FBM) if the memory process satisfies some conditions [5]. If the process 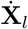 is covariance stationary and have the spectral density

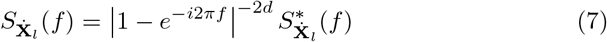

for 0 < *d* < 1/2, *X*, 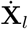 is said to have long memory, and 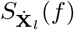 is called a *generalized spectral density* [26]. On the other hand, let 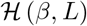 be the set of even non-negative functions *g* on [−1/2,1/2] such that

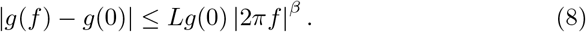

#### Theorem 1.

*Given the constants L,β such that* 0 < *L* < ∞, *β* ∈ (0, 2], *and* [*d_min_, d_max_*] ⊂ ((1 + *β*)/2 − *α, M* + 1/2), *there exists a constant C* > 0 *such that, for all j* ≥ 0, *d_l_* ∈ [*d_min_, d_max_*] *and* 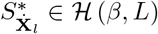,

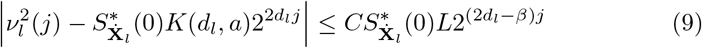

*where*

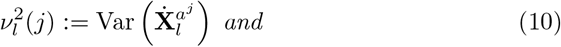

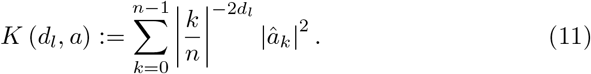

The theorem was proved by Moulines *et al*. in [5]. They showed that 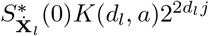 is the spectral density of the filtered series of a generalized FBM *B_(d)_* which is defined as a mean-zero Gaussian process with covariance

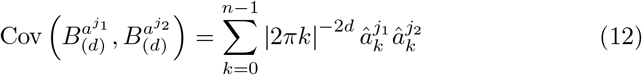

for *d* ∈ (1/2 – *α, M* + 1/2). Therefore, from (9), 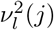 can be well approximated by 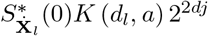; i.e.

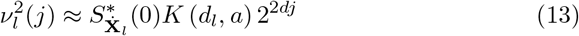

if *d_l_* < *β*/2. By denoting 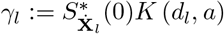, we have the following regression model

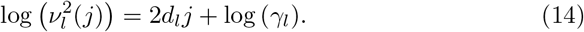

Since 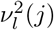 can be estimated by the empirical variance 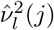, we can also obtain estimator of the memory parameter *d* through the ordinary regression estimator

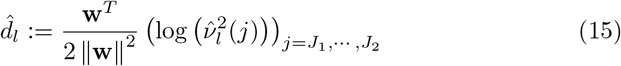

where the vector **w** := [*w*_0_, ⋯, *w_l_*]^*T*^ satisfies 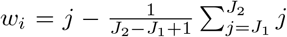. From (14) and (15), we have 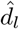 and 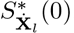.

This least squares estimator of long memory parameter is similar with that of fractional Brownian motion discussed by Coeurjolly and Achard [27][7]. For example, they described that a fractional Brownian motion **B**_*H*_ with Hurst parameter *H* ∈ (0,1) and scaling coefficient *C* > 0 has the following property

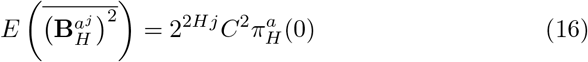

where 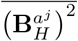 denotes the empirical mean of the squared filtered coefficients, and

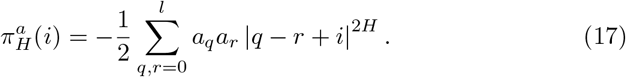

The equation (16) of a FBM is similar with the equation (13) of a long memory process, which indicates that Achard *et al*.’s method [7] for estimating the Hurst parameter of a fractional Brownian motion in the presence of additive noise is applicable to the general class of long memory processes without any modification. In the next sections 3.3 and 3.4, we will summarize how to apply Achard *et al*.’s method to general long memory processes.

### 3.3 Model B0: Long Memory Process with an Additive Brownian Motion

Let us consider the following model

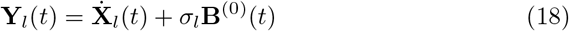

where *σ* > 0, 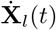 is a long memory process with parameter *d_l_* as defined in the section 3 and **B**^(0)^(*t*) for *t* =1, ⋯, *N* is a standard Brownian motion. Then, from (13), the variance of the filtered series of **Y**_*l*_ will be

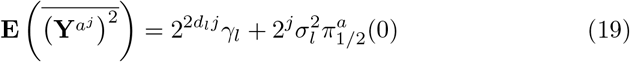

where 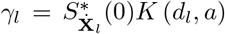. Thus, we can apply the same method as Achard *et al*. did as follows; if we define 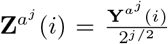, the least squares estimate is computed by

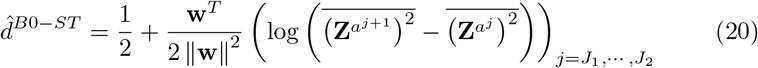

### 3.4 Model B1: Long Memory Process with an Additive White Noise

Let us consider the following model

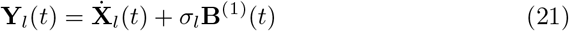

where *σ_l_* > 0 and **B**^(1)^(*t*) for *t* = 1, ⋯, *N* are i.i.d. standard Gaussian variables. According to Achard *et al*. [7],

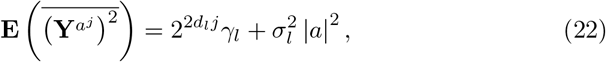

and the least squares estimate is

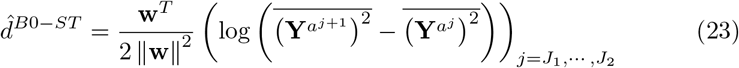

## 4 Linear Combination of Lone Memory Processes

In the previous section, we dealt with two models: a fractional Brownian motion contaminated by a standard Brownian motion, and a fractional Brownian motion contaminated by an additive white noise. Here, we will generalize the signal model; let us consider the following model

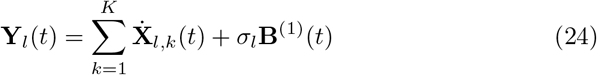

where *K* is the unknown integer, 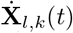 for *k* =1, ⋯, *K* is a long memory process with parameter *d_l,k_* and **B**^(1)^(*t*) for *t* = 1, ⋯, *N* are i.i.d. standard Gaussian variables. The variance of the filtered series of **Y**_*l*_ will be

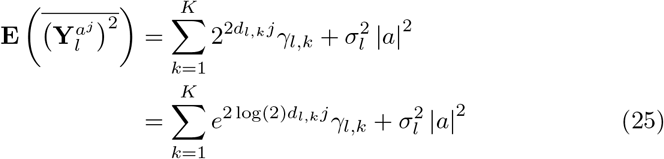

where 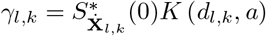. Then, we would like to estimate *K* and *d_l,k_* for all *k* = 1, ⋯, *K* given the measurement 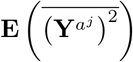 for *j* = *J*_1_, ⋯, *J*_2_.

This problem can be interpreted as the *multiexponential analysis* which is a common problem in the various fields such as physics, chemistry, medical imaging, and so forth. There exist numerous methods such as nonlinear least squares analysis (grid search, gradient search, Gauss-Newton method, Levenberg-Marquard method) [1] and Bayesian probability theory [2]. Even though in general the solution of this problem is not unique [1], it can be converted to the well-posed problem by applying some constraints on parameters.

The other issue is that the estimation is sensitive to the number of scales *J*_2_ – *J*_1_ + 1 and the resolution of parameters. In otherwords, the decrease in the number of scales and the increase in the number of components *K* or the resolution *d_l,k_1__*/*d_l,k_2__*(if *dl,k_1_* > *d_l,k_2__* for *k*_1_, *k*_2_ = 1, ⋯, *K*) make the parameter estimation less precise. In the fMRI analysis, the number of scales is normally less than 10 which is tremendously small.

To simplify our problem, let us consider the following model

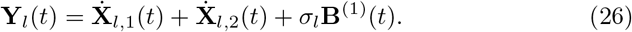

This model is similar with the original fMRI signal model defined in (1). If we successfully estimate the long memory parameters of the signals 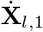, and 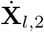, we will be able to classify them into endogenous signal **X**_*l*_ and physiological noises (such as **C**_*l*_ or **R**_*l*_). From (26), the variance of filtered series of **Y**_*l*_ is

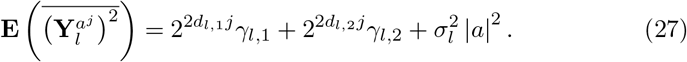

In the framework of the nonlinear least squares methods [1], our problem can be formulated as finding out the set of optimal parameters (*d*_*l*,1_, *d*_*l*,2_, *γ*_*l*,1_, *γ*_*l*,2_, 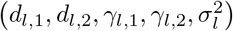) which minimizes the least squares error 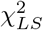 such that

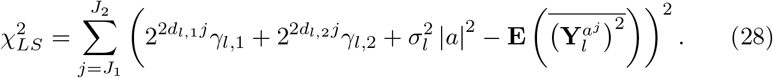

To solve the problem, we exploited the following simple algorithm. Let 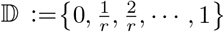 be a discrete set of long memory parameters where *r* is the resolution of long memory parameter. Then, we computed the following quantity for all 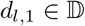 and 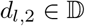

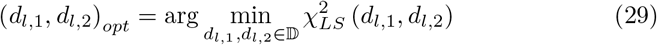

where

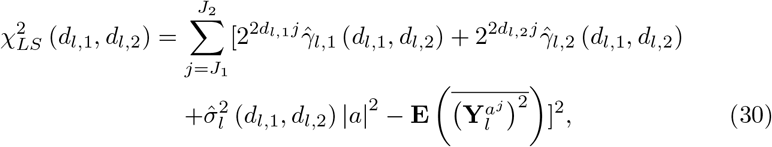

and the vector 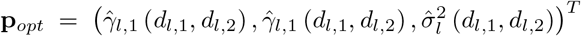 is determined by

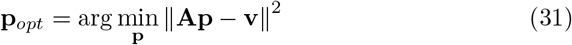

where

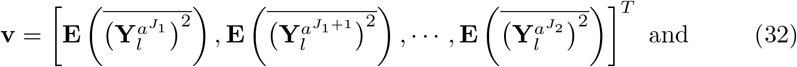

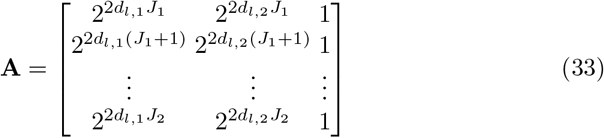

given *d*_*l*,1_ and *d*_*l*,2_ [8].

## 5 Experimental Results

### 5.1 Long Memory Processes Corrupted by White Noise

To demonstrate that Achard *et al*.’s method [7] can be directly applied to general long memory processes, we simulated ARFIMA processes [9] contaminated by additive white noise since the process belongs to the class of long memory processes [6]. The process **X** is called the ARFIMA(*p, d, q*) process if *S*\*(*f*) is given for −1/2 < *d* < 1/2 by

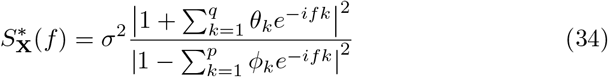

with 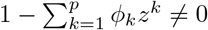 for |*z*| = 1 [6].

To estimate the long memory parameter of the ARFIMA processes, we exploited the R-package `dvfBm` available on the `R CRAN` (http://cran.r-project.org). We will use the same notation as Achard *et al*. defined in [7]; that is, 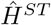 is a standard least squares estimator, 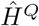 is a least squares estimator based on sample quantiles, and 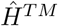 is a least squares estimator based on trimmed mean. Also, 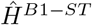, 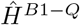, and 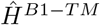 denote least squares estimators when the time series are assumed to be contaminated by white noise. Notice that the experimental results do not show the mean and standard deviation of the estimators since we did not run replication for each time series. We set up the specific parameters as follows: **p** =1/2 and **c** = 1 for 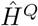, and *β*_1_ = *β*_2_ =0.1 for 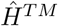. We only used the filter i1 which corresponds to the increments of order 1, and the minimum *M*_1_ and the maximum *M*_2_ of filter dilation were set up as 1 and 5. The whole results are shown at the tables 1, 2, 3, 4, 5, 6, 7, and 8.

**Table 1:** Long memory estimates of an ARFIMA (0, *d*, 0) process for *d* = 0.2, 0.4.

|  | d=0.2 |  |  | d=0.4 |  |  |
| --- | --- | --- | --- | --- | --- | --- |
|  | n=100 | n=1000 | n=10000 | n=100 | n=1000 | n=10000 |
| ST(i1,5) | 0.160 | 0.172 | 0.159 | 0.442 | 0.374 | 0.366 |
| Q(i1,5) | 0.266 | 0.155 | 0.159 | 0.512 | 0.349 | 0.365 |
| TM(i1,5) | 0.174 | 0.173 | 0.158 | 0.495 | 0.355 | 0.366 |

**Table 2:** Long memory estimates of an ARFIMA (1, *d*, 0) process with *ϕ*_1_ = −0.2 for *d* = 0.2, 0.4.

|  | d=0.2 |  |  | d=0.4 |  |  |
| --- | --- | --- | --- | --- | --- | --- |
|  | n=100 | n=1000 | n=10000 | n=100 | n=1000 | n=10000 |
| ST(i1,5) | 0.337 | 0.304 | 0.290 | 0.463 | 0.555 | 0.530 |
| Q(i1,5) | 0.397 | 0.283 | 0.280 | 0.402 | 0.550 | 0.527 |
| TM(i1,5) | 0.364 | 0.310 | 0.283 | 0.456 | 0.558 | 0.529 |

**Table 3:** Long memory estimates of an ARFIMA (0; *d*; 1) process with *θ*_1_ = −0.3 for *d* = 0.2, 0.4.

|  | d=0.2 |  |  | d=0.4 |  |  |
| --- | --- | --- | --- | --- | --- | --- |
|  | n=100 | n=1000 | n=10000 | n=100 | n=1000 | n=10000 |
| ST | 0.050 | 0.026 | 0.016 | 0.097 | 0.203 | 0.185 |
| Q | 0.105 | 0.031 | 0.015 | 0.208 | 0.210 | 0.179 |
| TM | 0.090 | 0.036 | 0.019 | 0.114 | 0.195 | 0.182 |

**Table 4:** Long memory estimates of an ARFIMA (10, *d*, 0) process with *ϕ_i_* = −0.2 for *d* = 0.2, 0.4 and *i* = 1, ⋯, 10.

|  | d=0.2 |  |  | d=0.4 |  |  |
| --- | --- | --- | --- | --- | --- | --- |
|  | n=100 | n=1000 | n=10000 | n=100 | n=1000 | n=10000 |
| ST | 1.718 | 1.686 |  | 1.718 | 1.686 |  |
| Q | 2.155 | 2.149 |  | 2.128 | 2.150 |  |
| TM | 1.971 | 1.942 |  | 1.971 | 1.942 |  |

**Table 5:**
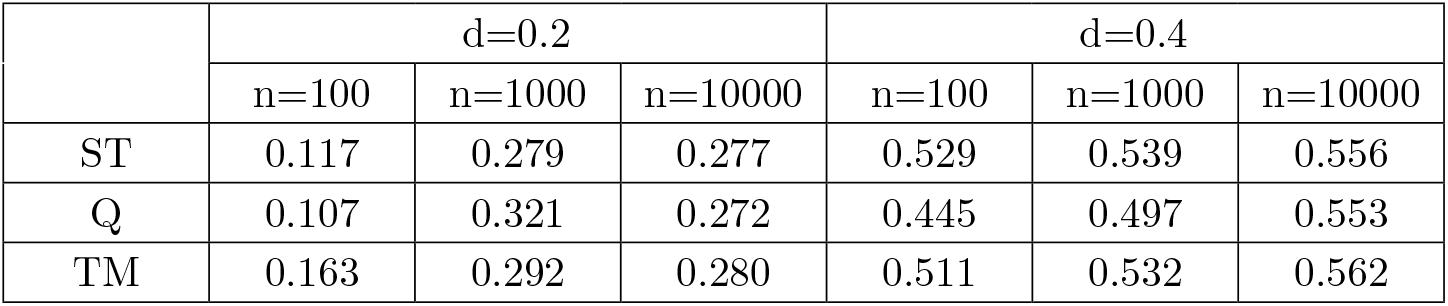
Long memory estimates of an ARFIMA (0, *d*, 10) process with *θ_i_* = −0.3 for *d* = 0.2, 0.4 and *i* = 1,⋯ 10.

**Table 6:** Long memory estimates of an ARFIMA (0, *d*, 0) process contaminated by white noise with a *SNR* = −10 for *d* = 0.2, 0.4.

|  | d=0.2 |  |  | d=0.4 |  |  |
| --- | --- | --- | --- | --- | --- | --- |
|  | n=100 | n=1000 | n=10000 | n=100 | n=1000 | n=10000 |
| ST | 0.097 | 0.035 | 0.022 | -0.041 | 0.041 | 0.052 |
| B1-ST | 0.049 | 0.297 | -0.182 | 0.271 | -0.360 | 0.148 |
| B1-Q | 0.303 | -0.497 | -0.283 | 0.319 | 0.354 | 0.083 |
| B1-TM | -0.032 | 0.238 | -0.143 | 0.212 | 0.472 | 0.086 |

**Table 7:** Long memory estimates of an ARFIMA (0, *d*, 0) process contaminated by white noise with a *SNR* = −20 for *d* = 0.2, 0.4.

|  | d=0.2 |  |  | d=0.4 |  |  |
| --- | --- | --- | --- | --- | --- | --- |
|  | n=100 | n=1000 | n=10000 | n=100 | n=1000 | n=10000 |
| ST | 0.015 | 0.021 | 0.007 | 0.035 | 0.002 | 0.005 |
| B1-ST | 0.056 | 0.004 | -0.255 | 0.014 | 0.415 | 0.539 |
| B1-Q | -0.095 | 0.330 | -0.326 | 0.213 | 0.430 | -0.225 |
| B1-TM | 0.106 | 0.330 | -0.546 | 0.336 | 0.683 | 0.324 |

**Table 8:** Long memory estimates of an ARFIMA (0, *d*, 10) process contaminated by white noise with a *SNR* = −20 and *θ_i_* = −0.3 for *d* = 0.2, 0.4 and *i* =1, ⋯, 10.

|  | d=0.2 |  |  | d=0.4 |  |  |
| --- | --- | --- | --- | --- | --- | --- |
|  | n=100 | n=1000 | n=10000 | n=100 | n=1000 | n=10000 |
| ST | 0.026 | -0.027 | 0.005 | 0.005 | 0.020 | 0.019 |
| B1-ST | -0.502 | 0.238 | 0.203 | -0.363 | -0.059 | 0.880 |
| B1-Q | -0.349 | -0.384 | 0.599 | -0.089 | 0.215 | 1.581 |
| B1-TM | -0.581 | -0.093 | 0.324 | -0.525 | -0.172 | 0.726 |

### 5.2 Linearly Summed fractional Brownian Motions

We tested the NLS long memory estimator of linearly summed fractional Brownian motions with the Hurst parameter *H*_1_ = 0.1, 0.2 and *H*_2_ = 0.6, 0.7, 0.8, 0.9. Each process was simulated by the method proposed by Abry and Sellan [10]. (The MATLAB function can be found at http://www.mathworks.com/access/helpdesk/help/toolbox/wavelet/wfbm.html) To compute wavelet variances, we exploited the maximum overlay discrete wavelet transform (MODWT) with the Daubechies filter of order 8. The results are shown at the table 9.

**Table 9:**
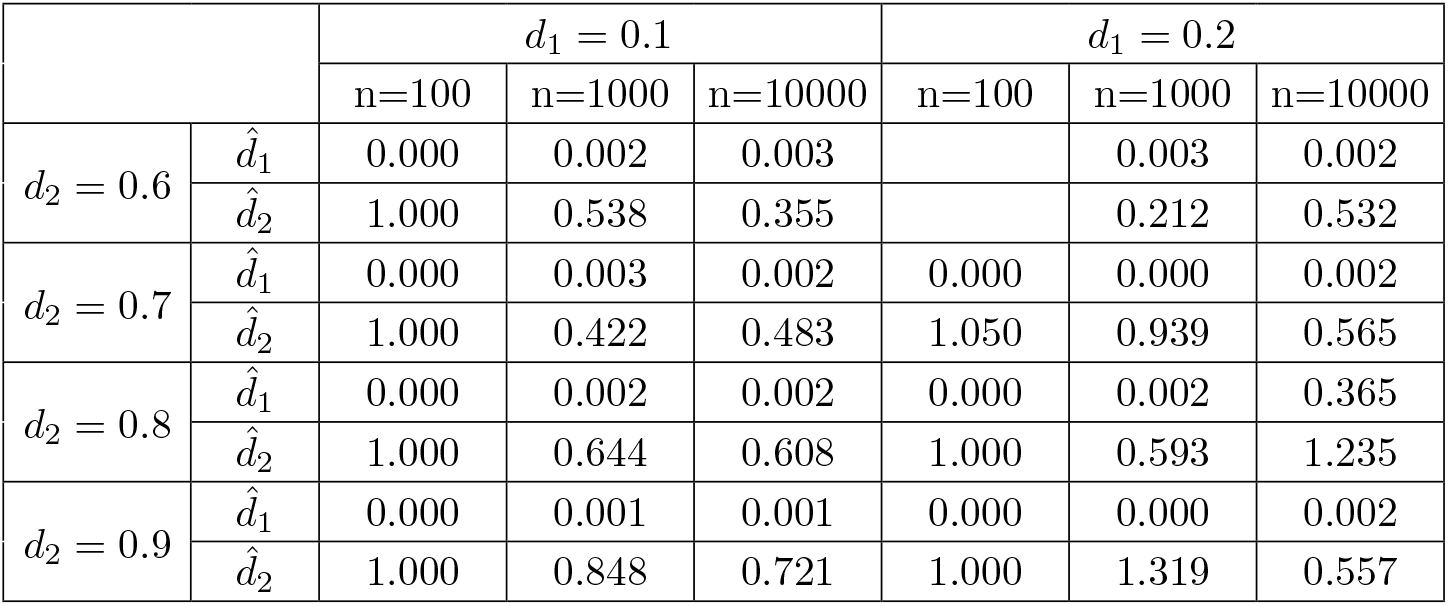
Long memory estimates of two linearly summed fractional Brownian motions for *H*_1_ = 0.1, 0.2, and *H*_2_ = 0.6, 0.7, 0.8, 0.9.

## 6 Discussion

### Estimation of long memory parameters in the presence of white noise

We applied Achard *et al*.’s method [7] to the ARFIMA(*p, d, q*) processes corrupted by white noises, but the performance was extremely poor except the case of ARFIMA(0, *d*, 0). We thought that the applicability of Achard *et al*.’s method to a long memory process **X** is based on the assumption such that 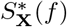 of the process **X** belongs to the function set 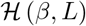 defined at (8). The fact that the parameter estimation of the ARFIMA(0, *d*, 10) process was better than that of the ARFIMA(0, *d*, 0) process implies that 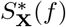 better fit into 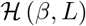 as the order *q* of MA part increases. Nevertheless, we need to understand exactly why the performance of estimation was significantly unacceptable, and to enhance Achard *et al*.’s method in order to make better performance for general long memory processes.

### Parameter estimation of linearly combined long memory processes

The parameter estimation of linearly combined long memory processes was seriously worse than that of a single long memory processes corrupted by white noise. As you see in the table 9, the estimates are not contiguous to the expected value. We need to verify how the NLS-based estimator is effective theoretically and empirically. As I noticed at the section 4, the small number of scales seems to contribute to the poor performance. In the future works, we will clarify what factors mostly influence the performance of estimation. In addition, we will attempt to develop the alternative methods for parameter estimation of linearly combined long memory processes.

## Footnotes

1 The notation of filter was adopted from Achard *et al*. [7], but its conditions are identical to those of the wavelet filter defined by Moulines *et al*. [5]. The only difference is that the conditions are based on Discrete Fourier Transform while Moulines *et al*.’s filter conditions are based on Continuous Fourier Transform. We just assume, without proof, that Moulines *et al*.’s theories are valid even in our conditions of the wavelet filter.

2 Notice that the notation *a^j^* is identical to *a*^2^*j*^^ in Achard *et al*.’s definition [7].

